# Loss of tumor suppressor WWOX accelerates pancreatic cancer development through promotion of TGFβ/BMP2 signaling

**DOI:** 10.1101/2022.03.10.483835

**Authors:** Hussam Husanie, Muhannad Abu-Remaileh, Kian Maroun, Lina Abu-Tair, Hazem Safadi, Karine Atlan, Talia Golan, Rami I. Aqeilan

**Affiliations:** The Concern Foundation Laboratories, The Lautenberg Center for Immunology and Cancer Research, Department of Immunology and Cancer Research-IMRIC, Faculty of Medicine, The Hebrew University of Jerusalem, Jerusalem, Israel; Department of Pathology, Hadassah Medical Center, Jerusalem, Israel; Oncology Institute, Sheba Medical Center, Tel Aviv University, Israel

**Keywords:** Pancreatic cancer, PDX, WWOX, KRAS, ADM, MAPK, Stat3, TGFβ, BMP2, SMAD3

## Abstract

Pancreatic cancer is one of the most lethal cancers, owing to its late diagnosis and resistance to chemotherapy. The tumor suppressor WW domain-containing oxidoreductase (WWOX), one of the most active fragile sites in the human genome (FRA16D), is commonly altered in pancreatic cancer. However, the direct contribution of WWOX loss to pancreatic cancer development and progression remains largely unknown. Here, we report that combined conditional deletion of *Wwox* and activation of *KRasG12D* in *Ptf1a-CreER*-expressing mice results in accelerated formation of precursor lesions and pancreatic carcinoma. At the molecular level, we found that WWOX physically interacts with SMAD3 and BMP2, which are known activators of the TGF-β signaling pathway. In the absence of WWOX, TGFβ/BMPs signaling was enhanced, leading to increased macrophage infiltration and enhanced cancer stemness. Finally, overexpression of WWOX in patient-derived xenografts led to diminished aggressiveness both *in vitro* and *in vivo*. Overall, our findings reveal an essential role of WWOX in pancreatic cancer development and progression and underscore its role as *a bona fide* tumor suppressor.

## Introduction

Pancreatic ductal adenocarcinoma (PDAC) is currently the fourth leading cause of cancer death in the modern world, with a five-year survival rate of 9% of the patients^1^. Approximately, 15-20% of patients are eligible for surgical resection, and the rest of the patients are inoperable or have advanced metastatic disease with minimal benefit from chemotherapy treatment^2^. PDAC is believed to develop gradually over time by accumulating mutations in several oncogenes and tumor suppressors. Constant activation of oncogenic *KRAS* is one of the most frequent mutations in human intraepithelial neoplasms (PanINs) and PDAC. For example, *Kras-G12D* was found to be a driver mutation in pancreatic mouse models needed for PanIN initiation, maintenance, and progression to PDAC^3^. Excessive *KRAS* activation has a wide range of outcomes in pancreatic acinar cells, including cell death, metaplasia, fibrosis development resembling chronic pancreatitis, and PDAC development. Nonetheless, mutations in tumor suppressor genes are also necessary for the progression of PanINs to PDAC, such as mutations in *CDKN2A*, *TP53* and *SMAD4*^4,5^. Despite the identification of a plethora of driver genes in PDAC, its prognosis remains the worst among human cancers^1^. Therefore, an in-depth understanding of the molecular mechanisms of additional genes associated with PDAC formation and the identification of new molecular targets and pathways involved in this process would enhance our understanding of PDAC initiation and progression.

Recent genome-wide exome sequencing studies identified new mutations and alterations that contribute to PDAC development^6^. Among these mutations, deletion of the WW domain-containing oxidoreductase (*WWOX*) gene is one of the most active and common fragile sites (CFSs) involved in cancer, has been identified^7^. WWOX has been mapped to the unstable PDAC subtype, which is characterized by genomic instability and defects in DNA maintenance^6^. Importantly, PDAC tumors harbor a reduction or even a loss of WWOX protein expression. All pancreatic cancer cell lines and 40% of primary tumors exhibit a significant reduction in WWOX protein expression^8^. Interestingly, WWOX expression has been shown to gradually decline with the progression of preneoplastic lesions^9^. In addition to genomic aberrations, hypermethylation of the WWOX promoter is evident in some pancreatic cancer cell lines and in primary pancreatic adenocarcinoma cases^8^. Furthermore, restoration of WWOX protein expression in pancreatic cancer cell lines induced apoptosis and suppressed tumorigenicity both *in vitro* and *in vivo*^8,10^. More recently, it has been reported that PDAC patients harboring a single nucleotide polymorphism (SNP) at *WWOX rs11644322 G>A* display worse prognosis and reduced gemcitabine sensitivity^10^. Despite numerous studies, the direct role of WWOX in PDAC initiation and progression remains largely unknown.

WWOX acts as an adaptor protein harboring two WW domains that interact with the proline-rich motifs of its partner protein, thereby regulating its stability and functions^11^. Both TGF-β and BMPs have been shown to promote PDAC^12,13^. It has been shown that WWOX physically binds to SMAD3 through its WW1 domain, leading to inhibition of its activity and enhanced TGF-β signaling in breast cancer cells^14^. In contrast, BMPs are conserved members of the TGFβ signaling pathway^15^, containing the PPXY motif, which is the binding motif of the WW1 domain of WWOX^16^. Nonetheless, the crosstalk between WWOX and BMPs has not yet been revealed.

In this study, we investigated the contribution of WWOX ablation in mice to PDAC initiation and progression and provided evidence that WWOX deletion leads to the acceleration of acinar to ductal metaplasia (ADM), PanIN lesion formation, and PDAC development by altering TGFβ/BMP signaling. We found that WWOX physically interacted with SMAD3 and BMP2, suppressing the activation of the TGF-β signaling pathway. Moreover, WWOX depletion was found to be associated with enhanced inflammation and the cancer stemness phenotype. Finally, restoration of WWOX in patient-derived xenografts (PDXs) of PDAC suppressed tumor growth. These findings indicate that WWOX loss accelerates PDAC initiation and formation by regulating TGFβ/BMP signaling.

## Results

### Targeted deletion of *Wwox* accelerates neoplastic lesions formation

To determine WWOX’s role of WWOX in PDAC formation, we generated a new genetically engineered mouse model harboring *Wwox* deletion and *Kras* activation in pancreatic acinar cells and examined their phenotypes. To achieve this, we bred mice carrying tamoxifen-inducible Cre (CreER) controlled by the *Ptf1a* specific acinar promoter (*Ptf1a-CreER*)^17^ with a conditional tdTomato reporter (*Rosa26*-LSL-*tdTomato*) to generate *Ptf1a-CreER; Rosa26-LSL-tdTomato* wild-type (WT) mice. These mice were then bred with conditional *Kras^G12D^* knock-in mice to generate *Kras+/LSL-G12D; Ptf1a-CreER; Rosa26-LSL-tdTomato* mice, hereafter referred to as KC mice. *Wwoxfloxed* mice (*Wwoxf/+* or *Wwoxf/f*)^18^ were bred with KC mice to generate KWC mice (*Wwox(f/f* or *+/f);Kras+/LSL-G12D; Ptf1a-CreER; Rosa26-LSL-tdTomato)* and WC mice (*Wwox(f/f)/(+/f); Ptf1a-CreER; Rosa26-LSL-tdTomato)* (Fig. 1A). The models were validated by detecting tomato expression in frozen sections of the tamoxifen-induced pancreas (Fig. S1A), immunoblot analysis (Fig. S1B) and WWOX immunohistochemical staining (Fig. S1C) and immunofluorescence staining for tdTomato and amylase (Fig. S1D). Successful tamoxifen activation of Cre was assessed in mice one month after post-tamoxifen injection by validation of tomato expression using immunofluorescence and specific acinar ablation of WWOX and pERK using immunohistochemical staining (Fig. 1B).

**Figure 1.**
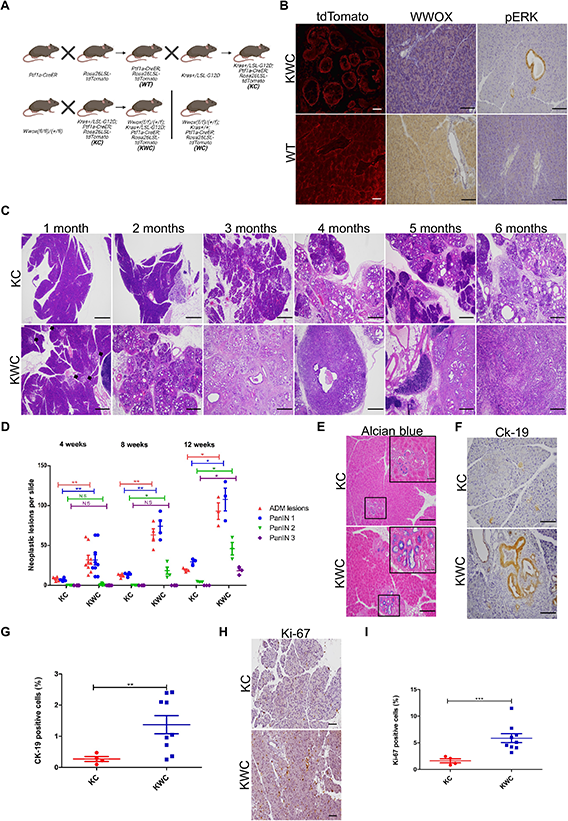
*Wwox* deletion synergies with *Kras^G12D^* activation by accelerating neoplastic lesions formation. (A) Illustration of KWC mice generation, *Ptf1a-CreER* mice were crossed with *Rosa26-LSL-tdTomato* to generate WT mice. WT mice were crossed with *Kras+/LSL-G12D* mice to generate KC mice. KC mice were crossed with *(Wwoxf/f)* mice to generate either KWC or WC mice. (B) Immunofluorescence of tdTomato (magnification, ×40), immunostaining of WWOX and pERK respectively in KWC, and WT mice (magnification, ×20). (C) Representative haematoxylin and eosin (H&E) stain of KC and KWC pancreata at 1,2,3,4,5, and 6 months respectively (magnification, ×4), n≥3 for each time point. (D) Quantification of ADM and PanIN lesions in KC and KWC mice in 4, 8, and 12-weeks post-tamoxifen injection. Number of KC mice, n=4 at one month, n=4 at 2 months, and n=3 at three months. Number of KWC mice, n=9 at one month, n=4 at 2 months, and n=3 at three months. (E) Representative Alcian blue stain of KC and KWC mice, one-month post-tamoxifen injection (magnification, ×40). (F) Representative Ck-19 staining of KC (n=4) and KWC (n=9) mice, one-month post-tamoxifen injection (magnification, ×20). (G) Quantification of Ck-19 in (Fig. 1F). (H) Representative Ki-67 staining of pancreata of KC (n=4) and KWC (n=9) mice, one-month post-tamoxifen injection (magnification, ×20). (I) Quantification of Ki-67 in (Fig. 1H). NS>0.05, \**P*<0.05, \*\**P*<0. 001, \*\*\**P*<0. 0001. Two-tail unpaired t-test. Quantification graphs are presenting the means ± SD.

Next, we compared the histological appearance of the pancreata between KWC and KC mice at different time points (1, 2, 3, 4, 5, and 6 months). WWOX deletion enhanced ADM lesions and PanIN formation in KWC mice compared to KC mice; lesions appeared as early as one month in KWC mice (Fig. 1C, black arrows). Moreover, some KWC mice started forming tumors as early as 4 months post tamoxifen injection, whereas none of the KC mice formed tumors at any of the experimental time points (Fig. 1C). However, no phenotypes were identified in the pancreata of WC mice, suggesting that *Wwox* deletion alone is insufficient for ADM and PanIN lesion formation (Fig. S1E). Quantification of ADM and PanIN lesions at 4, 8 and 12-weeks post-tamoxifen injection revealed a significant increase in the number in KWC mice compared to KC mice (Fig. 1D). Strikingly, no differences were observed between KWC mice harboring *Wwoxf/+* and *Wwoxf/f alleles*. Therefore, we refer to these mice as identical throughout our study.

Furthermore, Alcian blue staining revealed an increased number of ADM lesions in KWC mice compared to that in KC mice (Fig. 1E). Moreover, immunostaining of Ck-19 (a marker of ductal cells) revealed an increased number of ductal cells in KWC mice compared to KC mice (Fig.1F-G). Additionally, a higher number of proliferative cells was observed in KWC mice than in KC mice (Fig. 1H-I). Taken together, our analysis revealed the acceleration of preneoplastic lesions in the KWC mice.

To dissect cell-autonomous versus non-cell-autonomous neoplastic lesion formation, acinar cells were isolated from KC and KWC mice, then cells were cultured to assess their ability to transdifferentiate into ductal cells. Consistent with previous results, acinar cells isolated from KWC mice transdifferentiated faster to ductal cells, showing higher levels of CK-19 and lower levels of amylase, than those isolated from KC mice (Fig. S1F-H). Moreover, tdTomato-sorted cells from mice two-months post-tamoxifen injection revealed an induction in *the Runx1*, *Onecut2*, and *Foxq1* genes, which are associated with ADM reprogramming^19^ (Fig. S1I). These findings suggest that WWOX loss accelerates KRAS-mediated ADM and PanIN lesions likely by facilitating transdifferentiation of acinar cells into ductal cells.

### KRAS activation induces WWOX expression

It has been previously reported that WWOX levels are induced by stress conditions, such as DNA double-strand breaks (DSBs)^20^. Therefore, we set to determine whether WWOX levels increased following KRAS activation in normal non-transformed cells. To test this hypothesis, we induced KRAS expression both *in vitro* and *in vivo*, and monitored WWOX expression. Strikingly, when comparing WWOX protein expression *in vivo* in WT and KC mice at normal acinar sites, levels were found to increase following KRAS activation, as assessed by immunohistochemistry (IHC) staining at one- and two-weeks post-tamoxifen injection (Fig. 2A). Immunoblotting of WWOX in WT and KC mice one month after tamoxifen injection further confirmed induction of WWOX protein expression (Fig. 2B). Moreover, qRT-PCR analysis revealed an upregulation of *Wwox* RNA expression in KC mice from sorted tomato acinar cells at one-months post-tamoxifen injection compared to acinar cells sorted from WT mice (Fig. 2C).

**Figure 2.**
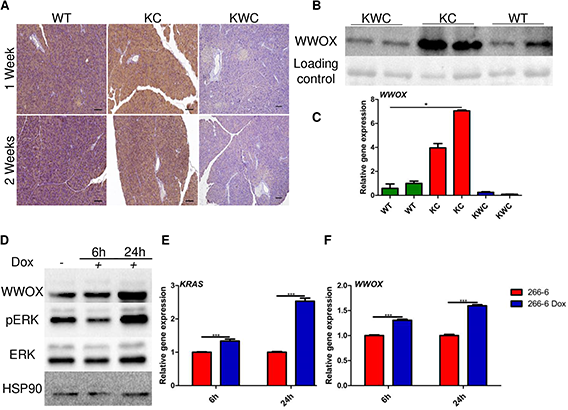
Induction of WWOX expression upon KRAS activation both *in vitro* and *in vivo*. (A) Representative WWOX immunostaining at normal sites of WT, KC and KWC mice one- and two-weeks post-tamoxifen injection. (B) Representative immunoblotting of WWOX of KWC (n=3), KC (n=3) and WT (n=3) mice one-month post-tamoxifen injection. (C) RT-PCR of *Wwox* in sorted acinar cells of WT, KC, and KWC mice one-months post-tamoxifen injection. (D) Immunoblotting of ERK, pERK, and WWOX in 266-6 cells activated/non-activated for 6h, and 24h. (E) RT-PCR of *Kras* in 266-6 cells activated/ non-activated for 6h, and 24h. (F) RT-PCR of *Wwox* in 266-6 cells activated/ non-activated for 6h, and 24h. \**P*<0.05, \*\**P*<0.001. Two-tail unpaired t-test. Quantification graphs are presenting the means ± SD.

These results are in contrast to those observed in patient samples, where WWOX expression has been shown to gradually decline upon PanIN formation^9^. To address this discrepancy, we examined WWOX protein expression in the pancreas of KC mice at different time points during lesion formation. Interestingly, we found that upon KRAS activation in KC mice, cytoplasmic WWOX expression gradually decreased with PanIN lesion progression (Fig. S2A-B). These results suggest differential expression of WWOX at short (1-2 weeks) vs. extended time points (1-2 months), indicating a tumorigenic barrier role of WWOX upon oncogenic KRAS activation, which is elevated immediately after RAS activation and then declines with lesion progression.

To further validate these results in an *in vitro* system, 266-6-Tet on/off KrasG12D murine acinar cells were treated with doxycycline for 6 and 24 h. Following 6h of activation, immunoblotting of pERK (a downstream target of KRAS) and WWOX revealed mild activation of KRAS and slight induction of WWOX, respectively (Fig. 2D). Remarkably, after 24h of KRAS activation, increased levels of pERK and WWOX were observed (Fig. 2D). Moreover, qRT-PCR analysis revealed upregulation of *Wwox* after KRAS activation for 6 and 24h (Fig. 2E-F). Taken together, these results suggest that WWOX is induced upon KRAS activation, perhaps as a response to reduced oncogenic stress.

### WWOX depletion enriches for EMT and cancer stem cells (CSCs)

To gain further insight into the molecular changes that accelerate lesion formation upon WWOX depletion, a comparative genome-wide transcriptional profiling was performed. Age-matched (two-months) WT (n=3), KC (n=3), and KWC (*Wwoxf/+* n=3 and *Wwoxf/f* n=2) cells from tomato-sorted acinar cells were used for bulk RNA sequencing (Supplementary Table 1). A set of 612 differentially regulated genes was identified between KWC and KC mice (Supplementary Table 2). Principal component analysis (PCA) showed a clear separation among all groups (WT, KC, and KWC mice) (Fig. 3A). The differentially expressed genes were enriched for pathways known to be implicated in WWOX signaling and have implications in PDAC (Fig. 3B) (Supplementary Table 3).

**Figure 3.**
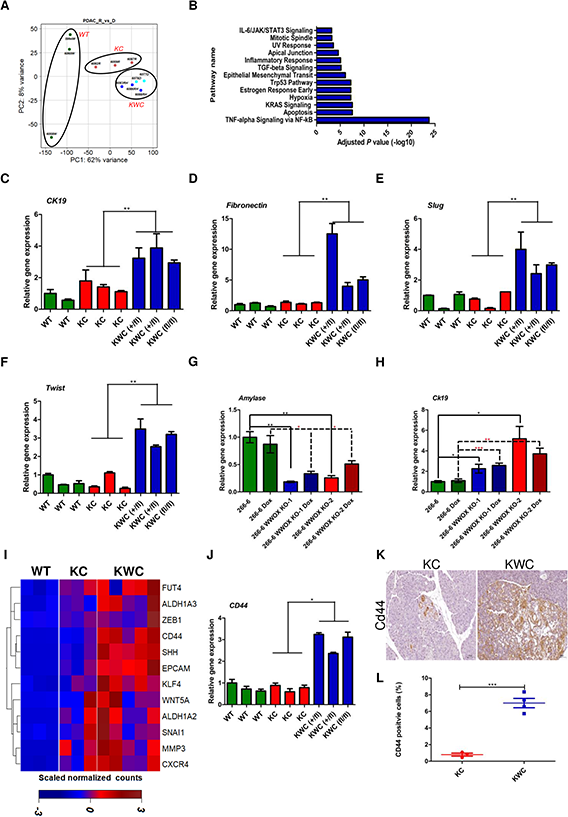
EMT and CSCs are accelerated in WWOX ablated pancreata. (A) PCA of WT mice (n=3), KC mice (n=3), KWC mice (n=5) two-months post-tamoxifen injection. (B) Molecular Signatures Database analysis (MSigDB) of differential expressed genes between KC, and KWC mice. (C-F) RT-PCR of *Ck-19*, *Fibronectin*, *Slug*, and *Twist* respectively of WT, KC, and KWC mice one-month post-tamoxifen injection. (G-H) RT-PCR of *Amylase*, and *Ck-19* respectively of 266-6, 266-6 WWOX KO-1, and 266-6 WWOX KO-2 with and without Dox. (I) Hierarchically clustered heat map illustration showing differential expression of CSCs associated genes in WT, KC and KWC mice from RNA-seq analysis. (J) RT-PCR of *CD44* in WT (n=3), KC (n=3) and KWC mice (n=3) one-month post-tamoxifen injection. \**P*<0.05, \*\**P*<0. 001. Two tail unpaired t test. Quantification graphs are presenting the means ± SD. Each column represents a mouse. SD is between technical replicates. (K) Representative IHC stain of CD44 in KC (n=3) and KWC mice (n=4) one-month post-tamoxifen injection (magnification, ×40). (L) Quantification of CD44 in KC and KWC mice one-month post-tamoxifen injection. \*\*\**P*<0. 0001. Two-tail unpaired t-test. Quantification graphs are presenting the means ± SD.

Pathway analysis of the RNA-seq data revealed upregulation of TGF-β signaling upon WWOX deletion (Fig 3B). Activation of the TGF-β signaling pathway is a common event in PDAC^21^. Furthermore, it has been reported that WWOX inhibits the TGF-β signaling pathway through direct interaction with SMAD3^14^. TGF-β suppresses the growth of many cell types through its SMAD-dependent pathway. Nonetheless, one aspect in which tumors benefit from TGF-β signaling is the activation of the EMT process through SMAD-dependent or SMAD-independent pathway^22^. EMT, a crucial step in early embryonic morphogenesis^23^, can also play an important role in many types of cancer, such as pancreatic cancer^24^. To validate the enrichment of the EMT pathway in our KWC model, qRT-PCR for *Ck-19*, *fibronectin*, *slug*, *Runx2*, and *Angptl4* of independent aged-matched RNA (two-months) of WT (n=3), KC (n=3), and KWC mice (n=3) revealed an upregulation of these markers in KWC mice compared to KC mice (Fig. S3A-E). In the same line, EMT markers (*Ck-19*, *fibronectin*, *Slug*, and *Twist* were upregulated in KWC mice (n=3) compared to KC mice (n=3) already in one-month post-tamoxifen injected mice (Fig. 3C-F). Moreover, knocking out WWOX from 266-6 reduced *amylase* expression and enhanced *Ck-19* expression (Fig. 3G-H). These findings indicate that WWOX loss, both *in vitro* and *in vivo*, induces a mesenchymal/ductal reprogramming phenotype, consistent with our previous findings (Fig. S1F-H).

Increasing evidence suggests that EMT acquisition results in cancer stem cell (CSC) characteristics that tend to invade tumors that are more resistant to therapies^23^. CSCs can self-renew; these cells exist in the tumor niche and account for the invasion of tumors^25^. Moreover, CSCs are a major cause of tumor recurrence in patients^26^. Interestingly, our RNA sequencing data revealed enrichment of stem cell-associated genes in KWC mice (Fig. 3I). To further validate our results, qRT-PCR analysis of CD44 levels in isolated acinar cells of age-matched (one-month) WT (n=3), KC (n=3), and KWC mice (n=3) revealed upregulation of CSCs genes in KWC mice compared with KC mice (Fig. 3J). Moreover, immunochemical staining of CD44 revealed a higher number of CD44 positive cells in KWC mice than in KC mice (Fig 3K-L). These results support a role for WWOX in the regulation of genes associated with cancer stemness.

### WWOX ablation enhances EMT and CSCs phenotype through promotion of TGFβ/BMP2 signaling

Growth factors such as TGFβ and BMPs are implicated in initiating the EMT process^27,28^. Gene set enrichment analysis (GSEA) of differentially expressed genes between KWC and KC mice identified hyperactivation of the TGF-β signaling pathway (Fig. 3B). To further strengthen our results, WT (n=2), KC (n=3), KWC *Wwoxf/+* (n=3), and KWC *Wwoxf/f* (n=2) mice one-month post-tamoxifen injection were sacrificed then acinar cells were isolated based on tdTomato expression. The expression of WWOX in these mice was validated using immunoblotting (Fig. S4A). Interestingly, WWOX deletion led to an increase in pSMAD2/3, as revealed by immunoblotting in KWC cells compared to that in KC cells (Fig. 4A). Furthermore, upregulation of *Angptl4*, a downstream target of TGFβ and SMAD3, was observed in KWC compared to KC cells isolated from mice two months after post-tamoxifen treatment (Fig. S4B). To further validate these results, a new cohort of mice was sacrificed one-month post-tamoxifen injection, acinar cells were isolated from WT (n=3), KC (n=3), KWC *Wwoxf/+* (n=2), and KWC *Wwoxf/f* (n=1) mice, and RNA was isolated. qRT-PCR analysis for *Serpine1* and *Angptl4*, downstream targets of SMAD3^14^, revealed their upregulation in KWC mice compared to KC mice (Fig. 4B-C). On the other hand, analysis of RNA-seq data for BMP2 revealed upregulation in KWC mice compared to KC mice (Fig. S4C). This finding was further substantiated by isolating Tomato positive cells from WT (n=2), KC (n=3), KWC *Wwoxf/+* (n=3), and KWC *Wwoxf/f* (n=2) mice one-month post-tamoxifen injection and immunoblotted for BMP2 and pSMAD1/5/9 (Fig. 4A). Interestingly, WWOX deletion induced BMP2 expression and its downstream target, pSMAD 1/5/9, in KWC mice compared to KC mice (Fig. 4A). Moreover, *Runx2*, a downstream target of BMP2^29^, was upregulated in KWC mice compared to KC mice one month after post-tamoxifen injection (Fig. 4D).

**Figure 4.**
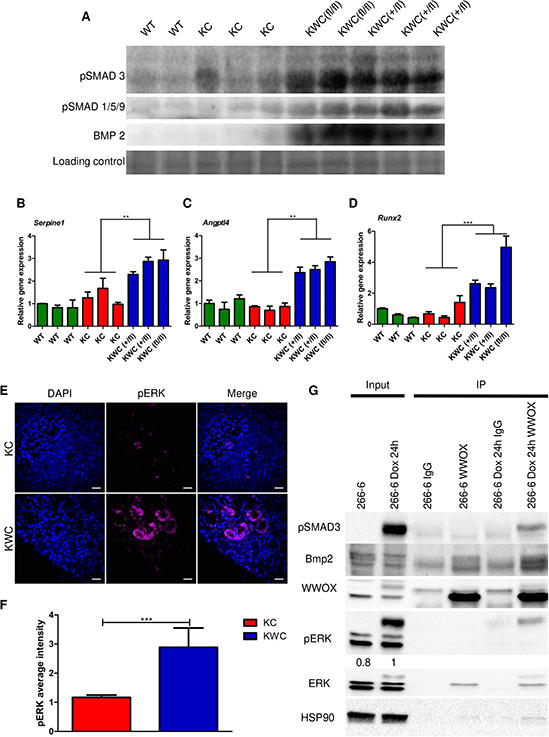
*Wwox* deletion promotes TGFβ and BMP2 through SMADs signaling. (A) Immunoblotting of pSMAD 2/3, pSMAD 1/5/9 and BMP2 of WT (n=2), KC (n=3), KWC *Wwoxf/+* (n=3), and KWC *Wwoxf/f* (n=2) mice one-month post-tamoxifen injection. (B-C) RT-PCR of *Serpine1* and *Angptl4* of WT, KC, and KWC mice one-month post-tamoxifen injection. (D) RT-PCR of *Runx2* of WT, KC and KWC mice one-month post-tamoxifen injection. (E) Representative immunofluorescence staining of pERK of KC, and KWC mice one -month post-tamoxifen injection (magnification, ×60). (F) Quantification of pERK staining of KC (n=4), and KWC (n=5). (G) Immunoprecipitation of endogenous WWOX from 266-6 cells not activated or activated (+dox) for 24 hours. Whole cell lysates were immunoprecipitated with either rabbit IgG (control) or monoclonal WWOX mouse antibodies. The immunoprecipitants were immunoblotted with anti-WWOX, anti-pSMAD2/3, anti-pERK, anti-ERK, and anti-BMP2/4 as indicated. \*\**P*<0. 001, \*\*\**P*<0. 0001. Two tail unpaired t test. Quantification graphs are presenting the means ± SD.

To check the possibility that WWOX deletion affects the ERK MAPK pathway, WT (n=2), KC (n=3), KWC *Wwoxf/+* (n=3), and *Wwoxf/f* (n=2) mice were sacrificed one month after post-tamoxifen injection, and acinar cells were isolated based on tdTomato expression and assessed by WWOX immunoblotting. Consistent with our previous results, WWOX deletion led to the induction of pERK in KWC mice compared to KC mice (Fig. S4A). Moreover, to validate the enrichment in KRAS signaling, pERK IHC staining in KC (n=4) and KWC (n=9) pancreata at one-month post-tamoxifen injection was performed. Strikingly, KRAS signaling was enhanced in KWC mice compared to that in KC mice at this early time point (Fig. S4D-E). To exclude the possibility that induction of KRAS signaling is due to the increased number of cells expressing KRAS because of the accelerated formation of preneoplastic lesions in KWC, immunofluorescence for pERK in KC (n=4) and KWC (n=5) pancreata at one-month post-tamoxifen injection was performed. Interestingly, pERK staining was more intense in KWC mice than KC mice (Fig. 4E-F). Altogether, these results imply that WWOX deletion enhances KRAS signaling through manipulation of TGF-β and BMP signaling.

### WWOX physically interacts with SMAD3 and BMP2 in acinar cells

WWOX acts as an adapter protein that regulates transcription, stability, and localization of its partners through physical interactions mediated by its WW1 domain^16^. For example, WWOX inhibits SMAD3 transcriptional activity via direct interaction in MCF10 cells^14^. To determine the occurrence of this interaction in pancreatic cells and whether WWOX interacts with the BMP family, we performed immunoprecipitation in 266-6 cells. To this end, inducible 266-6 KrasG12D murine acinar cells were activated for 24 h (+dox), and immunoprecipitation of WWOX was performed to check possible interactions with either SMAD3 and/or BMP2 proteins. Interestingly, we found that WWOX physically interacts with both SMAD3 and BMP2 following KRAS activation (Fig. 4G). These results further imply an important role for WWOX in regulating EMT and CSCs by regulating both the SMAD3 and BMP2 pathways upon KRAS activation.

### WWOX deficiency accelerates PDAC formation

Reduction or loss of WWOX protein expression has been associated with PDAC formation^9^.Furthermore, ablation of WWOX accelerated the formation of preneoplastic lesions (Fig.1). Taken together, these observations prompted us to further examine aging KC and KWC mice. To this end, mice were injected with tamoxifen at the age of one month, and then monitored and followed for another eight months. Interestingly, 16 out of 64 *Wwoxf/+* or *Wwoxf/f* KWC mice formed tumors between the ages of 4-8 months while none of the KC mice (n=14) formed tumors in this time range (Fig 5A), which is consistent with the literature^30,31^, as activation of KRAS alone is not sufficient for the formation of neoplastic lesions at these time points. To validate that the tumors obtained from KWC mice originated from *Ptf1a* positive cells, we stained the tumors with tdTomato antibody and detected them by immunofluorescence. We found that the tumors were positively stained for tdTomato, confirming that the acinar cells were the cell of origin. Moreover, histological characterization (H&E) of KWC tumors revealed highly aggressive tumors (Fig. 5B). Interestingly, *Wwoxf/+* KWC and *Wwoxf/f* KWC mice developed tumors at the same rate (Fig. 5A). Staining for WWOX protein expression in the tumors revealed that those developed in *Wwoxf/+* KWC mice displayed reduced or lost WWOX expression at the tumor sites compared to the normal sites, suggesting a loss of *Wwox* heterozygosity at these sites (Fig. 5C). Furthermore, the tumors formed were rich in spindle-shaped cell morphology, positive for pancytokeratin markers, and negative for amylase staining, indicating a mesenchymal identity (Fig. 5C). To further validate tumor aggressiveness, we stained for pERK and pSTAT3, and found that the tumor sites were highly positive for these markers (Fig. 5C). Moreover, these tumors were highly positive for vimentin and weakly positive for E-cadherin (a marker of EMT), which may indicate their ability to metastasize (Fig. 5D). Collectively, these data suggest the aggressiveness of KWC PDAC tumors.

**Figure 5.**
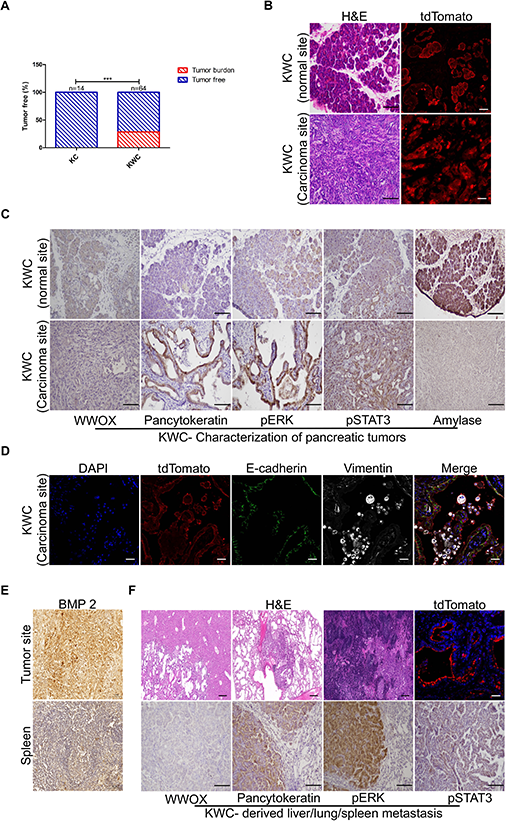
WWOX ablation accelerates PDAC formation in mice. (A) 16 out of 64 *Wwoxf/+* or *Wwoxf/f* KWC mice formed tumors between the ages 4-8 months while none of the KC mice (n=14) did in this time range. (B). Left panel-representative immunofluorescence stain of tdTomato of normal and carcinoma site of KWC mice (magnification, ×40). (C) Representative WWOX, Pancytokeratin, pERK, pStat3, and Amylase immunochemical stain of KWC mice at normal and carcinoma sites (magnification, ×10). (D) Representative immunofluorescence stain of tdTomato, E-cadherin, and Vimentin of KWC mice at carcinoma site (magnification, ×40). (E) Representative IHC staining of BMP2 at tumor site of a mouse at age of 8 months, and at spleen site of the same mouse (magnification, ×40). (F) Upper panel-H&E stain of liver, lung, and spleen (magnification, ×10). Representative immunofluorescence stain of tdTomato of metastasis site in the liver (magnification, ×40). Lower panel-Representative WWOX, Pancytokeratin, pERK, pSTAT3, and amylase immunochemical staining of metastasis site in the lung of KWC mice (magnification, ×10). \*\*\**P*<0. 0001. Two tail unpaired t-test. Quantification graphs are presenting the means ± SD.

### Enhanced progression and oncogenic signaling in KWC tumors

It has been demonstrated that some BMPs components, such as BMP2, are upregulated in pancreatic cancer patients compared to normal pancreatic tissue^32^. Moreover, our results indicate an enhancement in BMP2 signaling upon WWOX ablation (Fig. 4A and S4C). IHC staining of KWC tumors for BMP2 indicated the induction of its expression compared to that in the adjacent spleen (non-metastatic) from the same mouse (Fig. 5E).

PDAC tumors have a high capacity to metastasize to the adjacent organs. Common sites of metastasis include the liver, peritoneum, spleen, and lungs^33^. Moreover, metastasis requires PDAC cells to switch their epithelial characterization to a mesenchymal state^34^, a phenomenon that was observed in KWC mice tumors (Fig. 5D). Therefore, we examined these organs in tumor-bearing KWC mice. Liver, lung, and spleen metastases were found in three of the 16 KWC tumor-bearing mice (Fig. 5F, S5A). To validate the origin of these metastatic lesions, they were stained for tdTomato and found to be positive for the tomato reporter, validating PDAC as the origin of metastatic lesions (Fig. 5F). Moreover, these metastatic lesions were negative for WWOX and highly expressed pancytokeratin, pERK, and pSTAT3, as assessed by IHC staining, further validating that PDAC tumors were the origin of these metastatic lesions (Fig. 5F). In contrast, normal KWC lungs were positive for WWOX and negative for pancytokeratin, pERK, and pSTAT3 by IHC staining (Fig. S5B).

To exclude the possibility of *Ptf1a* promoter leakage to distant sites, such as the liver and lung, we stained WT mice expressing tomato after post-tamoxifen injection. As expected, the liver and lung tissues (n=3) were negative for tomatoes, whereas the pancreatic tissues were positive (Fig. S5C), suggesting that the primary tumor was the origin of metastases. Collectively, these data suggest the aggressiveness of KWC PDAC tumors and their ability to metastasize to the adjacent organs.

### WWOX deficiency in PDAC patients

To assess the value of WWOX expression in PDAC^8^, KM plotter dataset (www.kmplot.com) was used. We found that patients with low WWOX expression exhibited lower overall survival than those with high WWOX expression (Fig. 6A). To further validate the effects of WWOX using a human cell culture system, we assessed its expression in five PDX cell lines derived from advanced tumors^35^ (#50, 92, 114, 134, and 139). Immunoblotting of WWOX for the five PDX lines revealed a reduction in three of them relative to each other [#92, 134 and 139] (Fig 6B). Transduction of all PDXs using a lenti-WWOX vector or empty vector (EV) was then performed to generate stable lines, which were validated by immunoblotting for WWOX (Fig. 6B-C). To validate whether WWOX restoration had any biological effects *in vitro*, colony formation assays were performed for all lines (WT, EV, and WWOX overexpression (OE)). Interestingly, WWOX restoration in PDX92, 134, and 139 cells harbouring low WWOX expression successfully suppressed colony formation (Fig. 6D and Fig. S6A-B). In contrast, WWOX restoration in PDX50 or 114, harboring high WWOX expression had no effect on survival (Fig. S6C-D).

**Figure 6.**
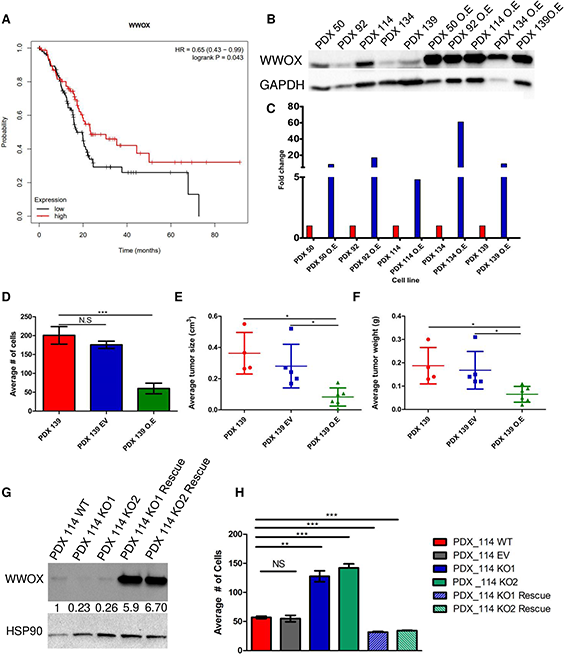
WWOX loss in human PDAC patients and PDX models. (A) Low expression of WWOX correlates with worse overall survival of PDAC patients as assessed using KMplotter dataset. The red line indicates high expression, while the black line low expression of WWOX. (B) Immunoblotting of WWOX in PDXs and PDXs-WWOX overexpression. (C) Quantification of WWOX expression from (B). (D) Quantification of colony formation of PDX-139, EV, and WWOX O.E, respectively. (E) Average tumor size of PDX-139 WT (n=4), PDX-139 EV (n=5), and PDX-139 WWOX O.E (n=6). (F) Average tumor weight of PDX-139 WT (n=4), PDX-139 EV (n=5), and PDX-139 WWOX O.E (n=6). (G) Immunoblotting of WWOX in PDX114, KO and KO rescue. (H) Quantification of colony formation of PDX114, EV, KO and KO rescue. NS>0.05, \**P*<0.05, \*\**P*<0. 001, \*\*\**P*<0. 0001. Two-tail unpaired t-test. Quantification graphs are presenting the means ± SD.

Next, we validated the effect of WWOX restoration on growth suppression *in vivo* using the PDX139 line. PDX139 WT, EV, and WWOX OE were intraperitonially injected into NOD/SCID mice. Indeed, PDX139 with WWOX overexpression resulted in smaller and lighter tumors compared to PDX139 WT and PDX139 EV (Fig. 6E-F), further confirming WWOX tumor suppressor activity. Staining of these tumors revealed higher expression of pERK in PDX139 and PDX139 EV than in PDX139 WWOX cells (Fig. S6E), consistent with our results in KWC mouse model.

To further examine the tumor suppressor function of WWOX in human PDXs, we used CRISPR/Cas9 to knockout WWOX in PDX114, which expresses high levels of WWOX, and assessed its effects on colony formation. PDX114 was transduced using either LentiCRISPR-V2 plasmid targeting WWOX exon 1 or re-expression of WWOX using CRISPR-untargetable WWOX mutant and generated stable polyclonal lines, which were validated by immunoblotting (Fig. 6G). Interestingly, both KO1 and KO2 clones have enhanced survival ability while restoring WWOX in these KO clones rescued this trait (Fig. 6H). It should be noted that rescue experiment led to supraphysiological levels of WWOX and this resulted in reduced survival, a hallmark of WWOX anti-tumor activity. Altogether, these findings further indicated the role of WWOX in tumor suppression in human pancreatic cancer cells.

## Discussion

Our current study demonstrates that somatic WWOX loss combined with oncogenic activation (*RasG12D*) in the Ptf1a mouse model resulted in accelerated formation of ADM, PanINs, and pancreatic tumors. Dissection of the plausible mechanism revealed that WWOX negatively regulates TGFβ/BMP signaling, and its loss is associated with the induction of SMADs signaling, downstream effectors of TGFβ/BMP signaling, which in turn enhances the expression of EMT markers, and CSCs phenotype, leading to acceleration in the formation of PanINs and PDAC development. These results further underscore the tumor-suppressive function of WWOX in pancreatic carcinogenesis.

Pancreatic cancer patients have the lowest survival rates among cancer patients, mainly due to late diagnoses and the limited benefits of standard-of-care treatments^1,7^. Therefore, understanding the molecular changes and pathways that contribute to PDAC is critical for improving the survival rates. In previous studies, WWOX was reported to be lost in pancreatic cancer cell lines, either by deletion or promoter hypermethylation^8,36^. In addition, restoration of WWOX expression in these cell lines suppressed cell growth both *in vitro* and *in vivo* and induces apoptosis^8^. These observations prompted us to examine the direct role of WWOX in PDAC. Previous studies have shown a gradual loss of WWOX expression with the progression of preneoplastic lesions in pancreatic cancer^8^. Here, we report for the first time the successful recapitulation of this phenomenon in a mouse model harbouring Ras activation, one of the major hallmarks of pancreatic cancer. Our findings revealed that WWOX acts as a barrier to halt tumor progression. When WWOX is lost, tumors can progress, as evidenced by the increased proliferation and appearance of malignant and metastatic tumors in KWC mice. This barrier seems to be critical in early ADM lesions, as KC mice exhibited a significant reduction in WWOX levels following the formation of preneoplastic lesions. Furthermore, the combined targeted deletion of *Wwox* and KRAS activation enables ADM reprogramming, an important step in PDAC development^37^. Recently, several specific genes have been proposed to be associated with the ADM process in pancreatic carcinogenesis, including *Onecut2* and *Foxq1*^19^. Our results showed that these genes were upregulated in tomato-positive cells isolated from KWC mice, as revealed by RNA-seq data.

Little is known about WWOX’s contribution of WWOX in pancreatic cancer at the molecular level. One study suggested that WWOX suppresses pancreatic cancer by upregulating SMAD4 signaling in Panc1 cells^8^. Using our pancreatic cancer model, we found that WWOX loss results in the activation of several known pathways implicated in PDAC formation, some of which have been previously linked to WWOX function, such as hypoxia^38^, DDR^39^ and TGFβ^40^ signaling. Interestingly, we revealed a mechanism by which WWOX alters TGF-β signaling and modulates its activity on target genes. The effect of WWOX on TGF-β is likely through direct binding to its downstream target SMAD3, as shown using the 266-6 system. SMAD3, an activator of TGF-β signaling, contributes to PDAC by promoting the EMT pathway, and its upregulation in PDAC patient samples predicts poor prognosis^41^, suggesting that WWOX may have an overall inhibitory role in EMT. Consistent with this notion, it has been reported that WWOX can inhibit the TGFβ signaling pathway in breast cancer cells by directly binding to SMAD3^14^.Our findings from a mouse model suggest that ablation of WWOX promotes TGFβ signaling by hyperactivating SMAD3 signaling and its specific targets, such as *Angptl4* and *Serpine1*^14^.

Several studies have described the crosstalk between ERK and SMAD signaling^42,43^ showing that TGF-β can induce ERK signaling through the PI3K/Pak2 pathway^44^. In contrast, BMPs can signal through the non-canonical pathway via different kinases^45^; for example, BMP4 can upregulate Dusp9 to attenuate ERK signaling in mouse embryonic stem cells^46^. Our study revealed a novel physical interaction between WWOX and BMP2 using a 266-6 cell culture system. BMPs are conserved members of the TGF-β signaling pathway, which regulate several biological processes, such as proliferation, apoptosis, EMT, and metastasis^47^. Moreover, BMP2 mRNA levels were increased in patients with PDAC, and there was a significant correlation between BMP2 elevated expression and reduced survival rates in PDAC patients^32^. Our findings in a mouse model suggest that deletion of WWOX promotes BMPs signaling by inducing the BMP2/SMAD1/5/9 axis.

WWOX has recently been reported to play a role in regulating EMT in endometrial adenocarcinoma patients^48^. In the current study, we found that loss of WWOX leads to enhanced TGF-β and BMPs signaling pathways, which in turn could induce the EMT phenotype. Upon WWOX deletion, mesenchymal markers of EMT, such as fibronectin, which is known to be an effector of WWOX signaling^49^. Moreover, transcription factors that regulate EMT, such as Slug and snail^50^ were upregulated. Accumulating evidence suggests that EMT acquisition results in CSC characteristics^23^. Furthermore, aberrant expression of WWOX has been shown to be associated with CSC markers in infiltrating breast cancer^51^. Here, we report that WWOX deletion is directly associated with elevation of CSC markers, such as CD44.

Several studies have described the crosstalk between ERK and SMAD signaling^42,43^. These studies have shown that TGF-β can induce ERK signaling through the PI3K/Pak2 pathway^44^. In contrast, BMPs can signal through the non-canonical pathway via different kinases^45^; for example, BMP4 can upregulate Dusp9 to attenuate ERK signaling in mouse embryonic stem cells^46^. Consistent with this notion, we found that WWOX deletion enhanced ERK signaling. This hyperactivation of ERK might be explained because of TGF-β and BMPs signaling promotion, which can crosstalk with the MAPK pathway and enhance its activation. This phenomenon should be further studied to understand whether hyperactivation of MAPK signaling is directly correlated with WWOX deletion.

Overall, our work revealed that the gene product of FRA16D, which is believed to be gradually lost in PDAC, might affect the early stages of PDAC formation as early as ADM formation. Our results underscore the role of WWOX as a driver tumor suppressor in PDAC and suggest that its loss or somatic deletion promotes tumorigenesis. WWOX promotes TGF-β signaling by interacting with its downstream target SMAD3. Moreover, WWOX interacts with BMP2, a member of the TGF-β family, which is known to be enhanced in patients with PDAC and a predictor of poor prognosis. Therefore, our findings suggest an important therapeutic value. Future studies should examine whether the use of TGF-β and BMPs inhibitors in low or absent WWOX-expressing PDAC tumors would suppress pancreatic tumorigenesis.

## Materials and methods

### Mice

Mice were purchased from the Jackson Laboratory (stock #00790875, stock #01937876, and stock #00817977) and crossed to generate wild-type mice (*Ptf1a-CreER; Rosa26-LSL-tdTomato)*, KC mice (*Kras+/LSL-G12D; Ptf1a-CreER; Rosa26-LSL-tdTomato)*, and *Wwox-floxed (Wwoxf/f)* C57BL6/J; 129sv mixed genetic background mice^18^ were crossed with KC mice to generate KWC mice: (*Wwox(f/f)/(+/f);Kras+/LSL-G12D; Ptf1a-CreER; Rosa26-LSL-tdTomato)*. Tamoxifen, dissolved in corn oil (Sigma: C8267, Sitnyakovo Blvd, Bulgaria), was subcutaneously injected into mice (male and female) aged four weeks, and two doses of 400 mg/kg were injected 48 h apart. Following tamoxifen administration, mice were checked twice a week and sacrificed at the experimental time points, as shown in the results. All experiments involving mice were approved by the Hebrew University Institutional Animal Care and Use Committee [#MD-18-15468-5].

### PDX injection in mice

Patient-derived xenografts (PDX) (PDX 50, 92, 114, 134, and 139) were obtained from Dr. Talia Golan (Sheba Medical Center)^35^. 1×10^6^ cells of PDX139, EV, and WWOX overexpression (OE) cells (1 × 106 cells) were suspended in DMEM and injected intraperitoneally (IP) into 6-8 weeks old NOD/SCID immunocompromised mice to study the ability of these human cells to form tumors. Tumor volume and weight were measured at the time of collection, 35-days after IP injection. The mice were euthanized, and primary tumors were collected for hematoxylin and eosin (H&E) and immunohistochemical (IHC) staining.

### Cell culture and plasmids

266-6 cells were obtained from Dr. Oren Parnas’s laboratory (Hebrew University of Jerusalem) and grown in DMEM (Gibco, San Diego, CA) supplemented with 10% FBS, 1% glutamine, 1% penicillin/streptomycin, and 1% sodium-pyruvate (Biological Industries, Beit-Haemek, Israel). Prior to culturing, plates were coated with 0.1% gelatin type A (Sigma: G2500, Sitnyakovo Blvd, Bulgaria) followed by refrigerated for 10-15 min, gelatin removed, and gently rinsed with medium before culturing the cells. For generating 266-6 KRASG12D cells, 266-6 CAS9 cells were infected with FUW-TetO-KRASG12D-BleoR virus and cells were selected with 5 mg/ml zeocin (Invitrogen, San Diego, CA) for 10 days. These cells were subsequently transduced with rtTA-tdTomato virus and were sorted for tdTomato positive cells by FACSAriaΙΙ. PDX cells^35^ were cultured in DMEM (Gibco, San Diego, CA) supplemented with 10% FBS, 1% glutamine, and 1% penicillin/streptomycin. To induce WWOX overexpression in PDX cell lines, cells were infected with a lentiviral vector containing wild-type WWOX and selected with 2 mg/mL G418 (Gibco:11811031, San Diego, CA). To induce KRAS expression in 266-6 cells 2 μg/mL of doxycycline (Tamar Laboratory Supplies) was added to the medium at various time points. All cells were grown at 37°C in 5% CO2 and frequently checked for mycoplasma.

### Colony formation assay

PDX cell line were plated at a density of 500 cells in 60 mm^2^ plates. After 10-14 days the cells were fixed with 70% ethanol, stained with Giemsa, and counted^52^.

### Histology and Immunohistochemistry

Tissues were fixed with 4% formalin, embedded in paraffin embedding and sectioning then stained with hematoxylin and eosin for histological. For immunohistochemistry (IHC), paraffin-embedded tissues were deparaffinized, followed by antigen retrieval using 25 mM citrate buffer (pH 6) in a pressure cooker. Then The sections were left to cool for 25 min, followed by blocking of the endogenous peroxidase with 3% H2O2 for 15 min. To reduce non-specific binding of the primary antibody, tissues were blocked with blocking solution (CAS Block) (Invitrogen, San Diego, CA), followed by incubation with the primary antibody overnight at 4°C. Sections were washed with TBST and incubated with secondary anti-rabbit, anti-mouse, or anti-rat immunoglobulin antibody for 30 min. The reaction was then performed using a DAB peroxidase kit (Vector Laboratories, SK-41 00, Mowry Ave Newark, United State), followed by hematoxylin stain for 45s as a counterstain. IHC stains were manually counted, at least five pictures were taken for each slide, and the average was calculated for each slide.

### Immunofluorescence stain from paraffin-embedded tissue

Paraffin-embedded sections were used for immunofluorescent (IF) staining. Deparaffinization and antigen retrieval were performed as previously described. sections were then blocked for one hour using a blocking solution containing 5% normal goat serum, 0.5% BSA, and 0.1% Triton X-100 in PBS (Biological Industries, Beit-Haemek, Israel). sections were then incubated with primary antibody at 4°C overnight. The following day, sections were washed three times with PBS and then incubated with the secondary antibody and Hoechst solution, diluted in blocking buffer, for one hour. Slides were washed three times with PBS, dried, and mounted with coverslips using fluorescence mounting medium (S3023; Dako, Santa Clara, United States). IF stains were manually counted, at least three confocal images were taken for each slide, and the average was calculated for each slide.

### Isolation of acinar cells and sorting

Mice were euthanized using CO_2_, followed by pancreas removal, and washing with cold HBSS (Biological Industries, Beit-Haemek, Israel). The pancreas was then cut into 1-2 mm pieces in cold HBSS and centrifuged at 600 rpm for 5 min at 4°C. For the first dissociation step, the pieces were resuspended in 0.05% trypsin-EDTA (solution C, Biological Industries, Beit-Haemek, Israel) at final concentration of 0.02% for 15 min shaking at 37°C, followed by washing with 10% FBS/DMEM to stop the reaction. The cells were centrifuged at 600 rpm followed by washing with 4% bovine serum albumin (BSA) (Mpbio, California, USA) prepared in HBSS containing 0.1 mg/ml DNase Ι (Sigma, Sitnyakovo Blvd, Bulgaria). For the second dissociation step, cells were resuspended with 4% BSA containing 0.1 mg/ml DNase Ι, and 10 mg/ml collagenase P (Sigma, Sitnyakovo Blvd, Bulgaria) at final concentration of 1 mg/ml of collagenase P. Cells were incubated for 20-30 min at 37°C with shaking, and they were pipetted up and down 10 times every 5 minutes of the incubation. cells were then passed through a 100 mm cell strainer (Corning, Massachusetts, USA) and washed with 4% BSA prepared in HBSS containing 0.1 mg/ml DNase Ι. The samples were then treated with red blood cell lysing buffer (Sigma Aldrich, Sitnyakovo Blvd, Bulgaria) and passed through a 40 μm cell strainer (Corning, Massachusetts, USA) and submitted for tdTomato sorting using BD FACSAria ΙΙΙ or Sony SH800.

### Seeding and culturing of acinar cells

After finishing from the digestion step from the isolation of acinar cells as mentioned before, cells were resuspended in 7 ml of Waymouth’s medium (Gibco, San Diego, CA) containing 2.5 % FBS, 1 % Penicillin-Streptomycin mixture, 0.25 mg/ml of trypsin inhibitor (Sigma, Sitnyakovo Blvd, Bulgaria), and 25 ng/ml of recombinant human Epidermal Growth Factor (EGF) (Sigma, Sitnyakovo Blvd, Bulgaria). Cells were then filtered through a 100 μm filter to retain the non-digested fragments, and then the cells were seeded in 6 well culture dish. Twenty-four hours after seeding, contaminating cells that adhered to 6 well cultured dish were eliminated, and cells in suspension were collected. The day before seeding the cells in suspension, a 6-well culture dish were coated with type I collagen (5 μg/cm2) (Biological Industries, Beit-Haemek, Israel), By adding 1 ml of type I collagen solution (50 μg/ml in 0.02 M acetic acid, 0.2 μm-filtered) to each well and allow it to passively adsorb on plastic, during 1 hr at 37 °C (or overnight at 4 °C). Collagen type I was aspirated, and the plates were allowed to dry under sterile conditions for 12 h. The acinar cells were then transferred to type I collagen plates. On the following day the medium was changed to eliminate non-viable cells that had not adhered, and the medium was changed each three days till day seven of the experiment. Cells were then stained for CK19 (Abcam; ab7755, Boston, USA) and amylase (Sigma; A8273, Sitnyakovo Blvd, Bulgaria) using immunofluorescence.

### RNA extraction, cDNA preparation and qRT-PCR

The TRI reagent (Biolab, Jerusalem, Israel) was used as described by the manufacturer to prepare the total RNA. To prepare cDNA, 1 μg of RNA was used with a QScript cDNA synthesis kit (Quantabio, Massachusetts, USA). The SYBR Green PCR Master Mix (Applied Biosystems, Massachusetts, USA) was used for qRT-PCR.

All measurements were performed in triplicate and normalized to the levels of *Ubc* or *Hprt* genes.

qRT-PCR was performed using the following murine primers sequence:

*Wwox_F*: 5′- GGGAGCTGCTACCACTGTCTA -3′.
*Wwox*_R: 5′- CCTCTCACTGAGTTCCCACA -3′.
*Ubc*_F: 5′- CAGCCGTATATCTTCCCA -3′.
*Ubc_R*: 5′- CTCAGAGGGATGCCAGTA -3′.
*Hprt*_F: 5′- TCAGTCAACGGGGGACATAAA -3′.
*Hprt*_R: 5′- GGGGCTGTACTGCTTAACCAG -3′.
*Snai2*_F: 5′- CGAACTGGACACACATACAGTG -3′.
*Snai2*_R: 5′- CTGAGGATCTCTGGTTGTGGT -3′.
*Serpine1_F*: 5′- GTCACCAACCGATTCGACCAG -3′.
*Serpine1*_R: 5′- GGGACTCTTTACGCAACTGTT -3′.
*Twist*_F: 5′- GTCCGCAGTCTTACGAGGAG -3′.
*Twist_R*: 5′- GCTTGAGGGTCTGAATCTTGCT -3′.
*Angplt4*_F: 5′- CATCCTGGGACGAGATGAACT -3′.
*Angptl4*_R: 5′- TGACAAGCGTTACCACAGGC -3′.
*Runx2*_F: 5′- CGGCCCTCCCTGAACTCT -3′.
*Runx2_R*: 5′- TGCCTGCCTGGGATCTGTA -3′.
*Ck-19_F*: 5′- GGGGGTTCAGTAATTGG -3′.
*Ck-19_R*: 5′- GAGGACGAGGTCACGAAGC -3′.
*Kras*_F: 5′- CAACAGTTGCGCAGCCTGAATGG -3′.
*Kras*_R: 5′- AAATCGCTGATTTGTGTAGTCGGT -3′.
*Fn1*_F: 5′- TACACCTGCTCCTGTGCTTC -3′.
*Fn1_R*: 5′-GAGACCTGCTCCTGTGCTTC -3′.
*Amylase*_F: 5′- CAGAGACATGGTGACAAGGTG -3′.
*Amylase*_R: 5′- ATCGTTAAAGTCCCAAGCAGA -3′.
*Cd-44*_F: 5′- TCGATTTGAATGTAACCTGCCG -3′.
*Cd-44_R*: 5′- CAGTCCGGGAGATACTGTAGC-3′.

### Immunoblotting

Total protein was lysed by using lysis buffer containing 50 mM Tris (pH7.5), 150 mM NaCl, 10% glycerol, 0.5% Nonidet P-40 (NP-40), with addition of protease and phosphatase inhibitors (1:100). The lysates were subjected to SDS-PAGE under standard conditions. The antibodies used were polyclonal WWOX^53^, monoclonal GAPDH (Calbiochem; CB1001, San Diego, USA), polyclonal HSP90 (Cell Signaling; 4874, Massachusetts, USA), monoclonal ERK (Cell Signaling; 4695, Massachusetts, USA), monoclonal pERK (Cell Signaling; 4370, Massachusetts, USA), polyclonal BMP2/4 (Santa Cruz; sc6267, Texas, USA), monoclonal pSMAD 2/3 (Cell Signaling; 8828, Massachusetts, USA), monoclonal pSMAD 1/5/9 (Cell Signaling; 13820, Massachusetts, USA).

### RNA-sequencing

Three WT, three KC, and five KWC male and female mice were euthanized using CO2 two months post-tamoxifen injection. Pancreata were disassociated and tdTomato acinar cells were sorted using BD FACSAria ΙΙΙ. Library preparation and sequencing were performed by the Genomic Applications Laboratory at Hebrew University’s Core Research Facility using a standard protocol. Briefly, RNA quality was checked using a tape station (Agilent Technologies; 5067-5576, California, USA), D1000 ScreenTape Kit (Agilent Technologies; 5067-5582, California, USA), Qubit(r) RNA HS Assay Kit (Invitrogen; Q32852, San Diego, CA), and Qubit(r) DNA HS Assay Kit (Invitrogen; 32854, San Diego, CA). For mRNA library preparation, 1 μg of RNA from each sample was processed using the KAPA Stranded mRNA-Seq Kit with mRNA capture beads (Kapa Biosystems, Bedford row, London). Each sample was then eluted in 20 μL of elution buffer adjusted to 10 mM, and 10 μL of each sample was collected and pooled in one tube. multiplex sample pool (1.5pM including PhiX 1.5%) was loaded in NextSeq 500/550 High Output v2 Kit (75 cycles) cartridge (Illumina; FC-404-1005) and loaded on NextSeq 500 System Machine (Illumina) with 75 cycles and single-read sequencing conditions. For library quality control, measures were applied to the raw FASTQ files, trimming low-quality bases (Q>20), residual adapters, and filtering out short reads (L>20). These measurements were performed using the Trim-Galore (v0.4.1) wrapper tool. File assessment before and after quality control was performed using FastQC (v0.11.5). The mouse transcriptome was mapped using salmon v0.13.1. Differential gene expression for the 11 samples was explored using the R package (v1.16.1) and analyzed using DESeq2 (v1.28.1).

### WWOX immunoprecipitation

266-6 cells were seeded in 14 cm plates. Cells were induced using doxycycline. 24 h, the cells were lysed using lysis buffer containing 50 mM Tris (pH7.5), 150 mM NaCl, 10% glycerol, 1% NP-40, protease, and phosphatase inhibitors. The samples were then subjected to a pre-cleaning step using mouse anti-IgG antibodies (Invitrogen, San Diego, CA). Following precleaning, samples were immunoprecipitated using monoclonal mouse anti-WWOX cocktail antibodies^53^ together with protein A/G plus agarose beads (Santa Cruz, Texas, USA) and incubated overnight at 4°C on a rotor. The next day beads were washed two times with lysis buffer, followed by three washes with washing buffer (NaCl 150nM, Tris 50 nM, Glycerol 5%, NP-40 0.05%, pH7.4).

### Targeting WWOX expression by CRISPR/CAS9

WWOX expression was knocked out using the LentiCRISPR-V2 plasmid (Addgene plasmid # 52961, Massachusetts, USA), which contains the sgRNA sequence that targets WWOX exon 1 [5’-CACCGCATGGCAGCGCTGCGCTACG-3’]^52^. Reexpression of WWOX using CRISPR-untargetable WWOX mutant was used in rescue experiments. Polyclonal pools were used in the experiments, and the cells were infected with an empty vector-V2 plasmid as a control.

## Acknowledgements

We are grateful to all members of the Aqeilan lab for fruitful discussions and suggestions. Special thanks to Mr. Jonathan Monin for assistance with bioinformatic analysis.

