## Supplementary for "Loss of tumor suppressor WWOX accelerates pancreatic cancer development through promotion of TGFβ/BMP2 signaling"

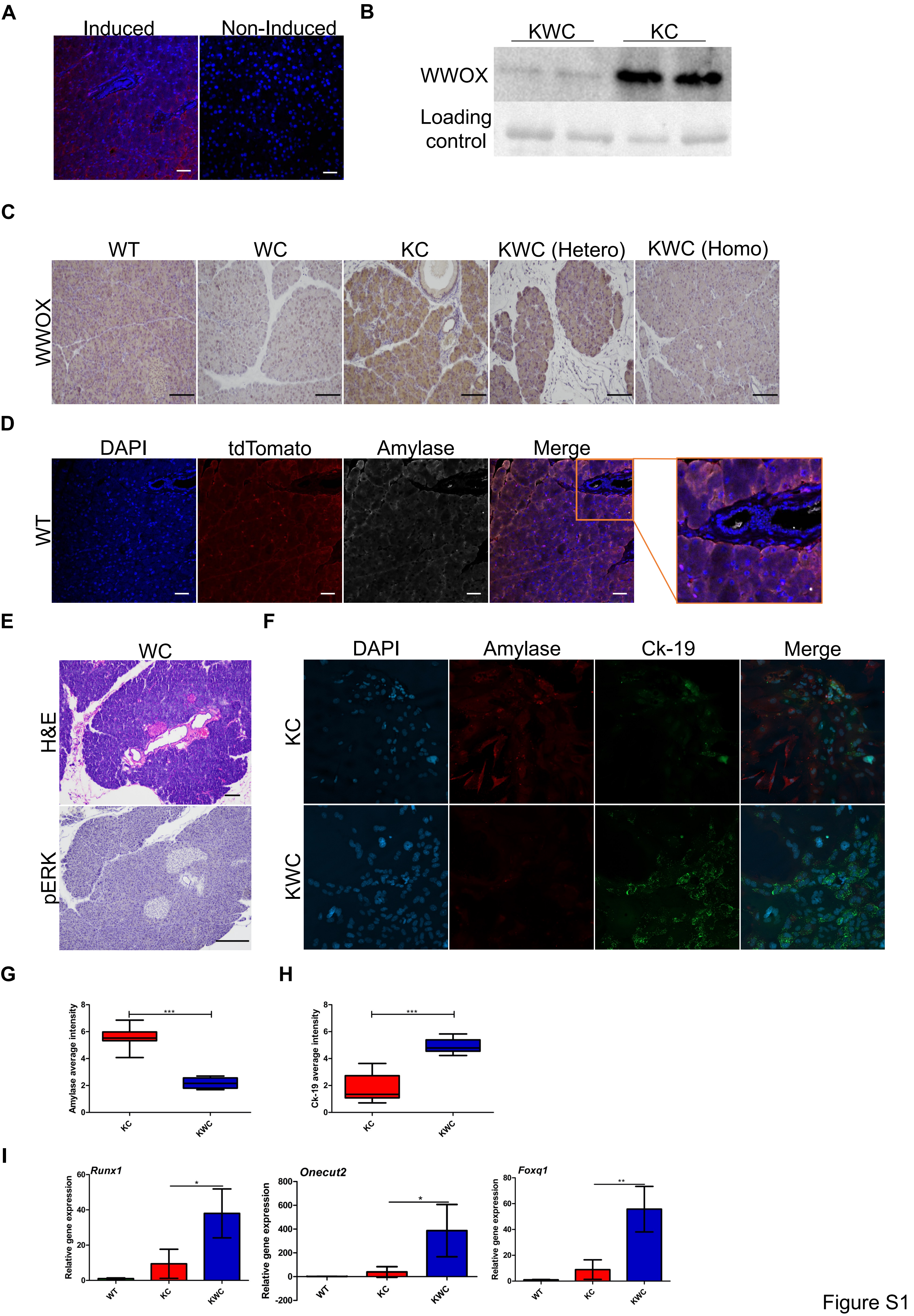
A
Induced
Non-Induced
B
KWC
KC
WWOX
Loading control
C
WT
WC
KC
KWC (Hetero)
KWC (Homo)
WWOX
D
DAPI
tdTomato
Amylase
Merge
WT
E
WC
H&E
pERK
F
DAPI
Amylase
Ck-19
Merge
KC
KWC
G
Amylase average intensity
***
KC
KWC
H
Ck-19 average intensity
***
KC
KWC
I
Runx1
Onecut2
Foxq1
Relative gene expression
*
**
WT
KC
KWC

Figure S1

**A**

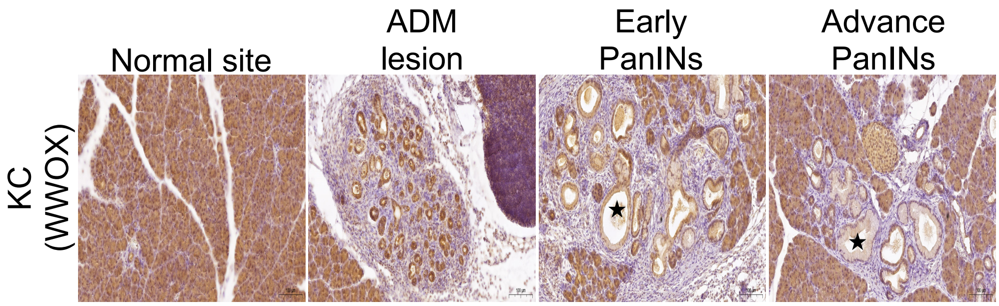
Normal site
ADM lesion
Early PanINs
Advance PanINs
KC (WWOX)

**B**

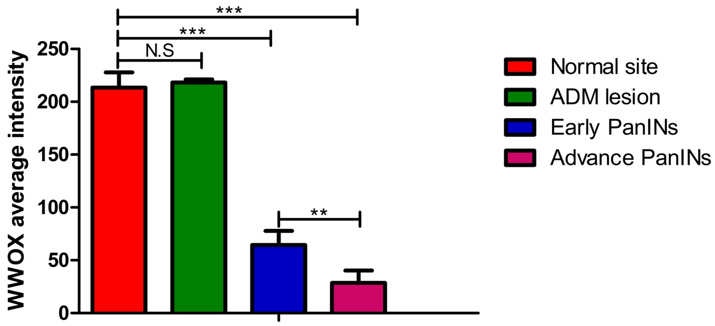
WWOX average intensity
250
200
150
100
50
0
***
***
N.S
**
Normal site
ADM lesion
Early PanINs
Advance PanINs

Figure S2

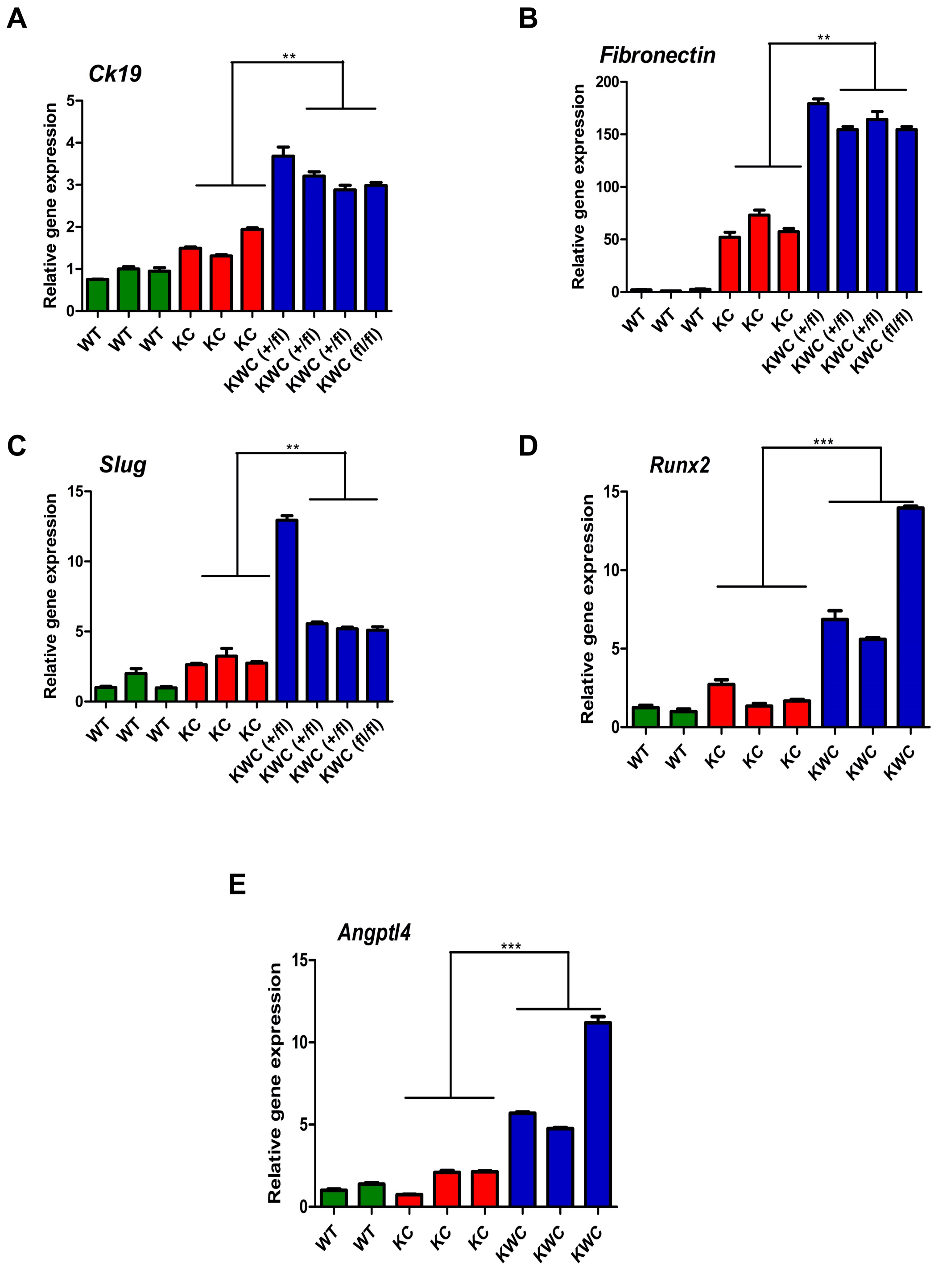

A
Ck19
Relative gene expression
0
1
2
3
4
5
**
WT
WT
WT
KC
KC
KC
KWC (+/fl)
KWC (+/fl)
KWC (+/fl)
KWC (fl/fl)
B
Fibronectin
Relative gene expression
0
50
100
150
200
**
WT
WT
WT
KC
KC
KC
KWC (+/fl)
KWC (+/fl)
KWC (+/fl)
KWC (fl/fl)
C
Slug
Relative gene expression
0
5
10
15
**
WT
WT
WT
KC
KC
KC
KWC (+/fl)
KWC (+/fl)
KWC (+/fl)
KWC (fl/fl)
D
Runx2
Relative gene expression
0
5
10
15
***
WT
WT
KC
KC
KC
KWC
KWC
KWC
E
Angptl4
Relative gene expression
0
5
10
15
***
WT
WT
KC
KC
KC
KWC
KWC
KWC


Figure S3

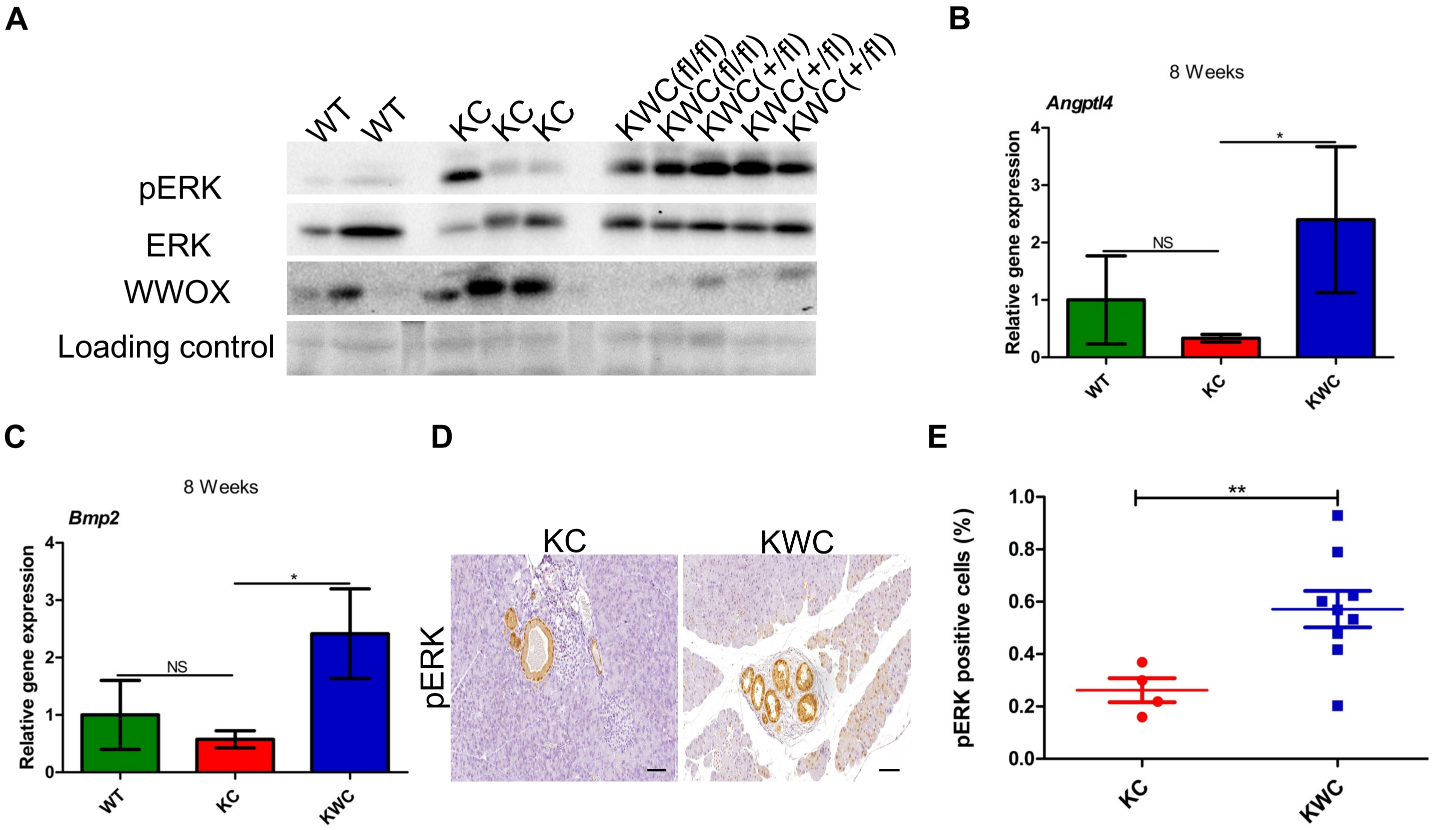
A
WT WT KC KC KC KWC(fl/fl) KWC(fl/fl) KWC(+/fl) KWC(+/fl) KWC(+/fl)
pERK
ERK
WWOX
Loading control
B
8 Weeks
Angptl4
Relative gene expression
NS
*
WT KC KWC
C
8 Weeks
Bmp2
Relative gene expression
NS
*
WT KC KWC
D
KC KWC
pERK
E
pERK positive cells (%)
**
KC KWC

Figure S4

**A**

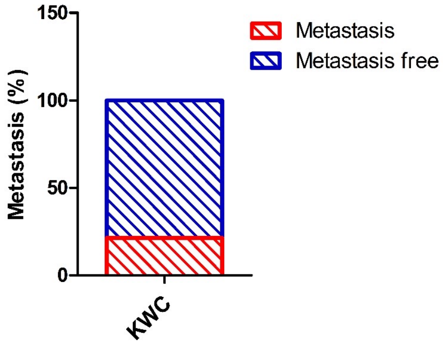
150
100
50
0
Metastasis (%)
Metastasis
Metastasis free
KWC

**B**

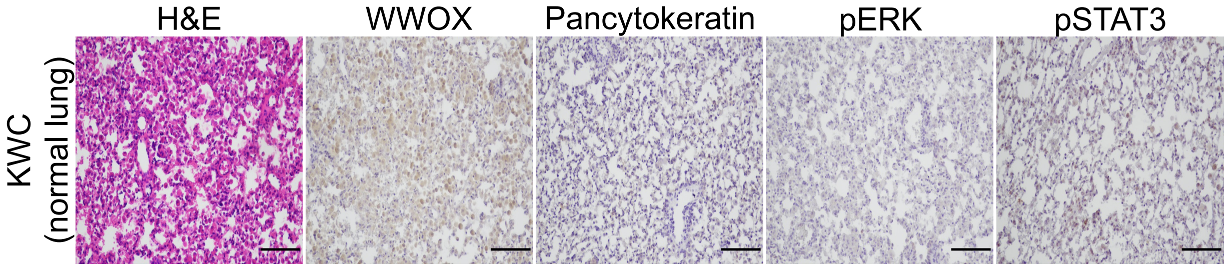
H&E
WWOX
Pancytokeratin
pERK
pSTAT3
KWC
(normal lung)

**C**

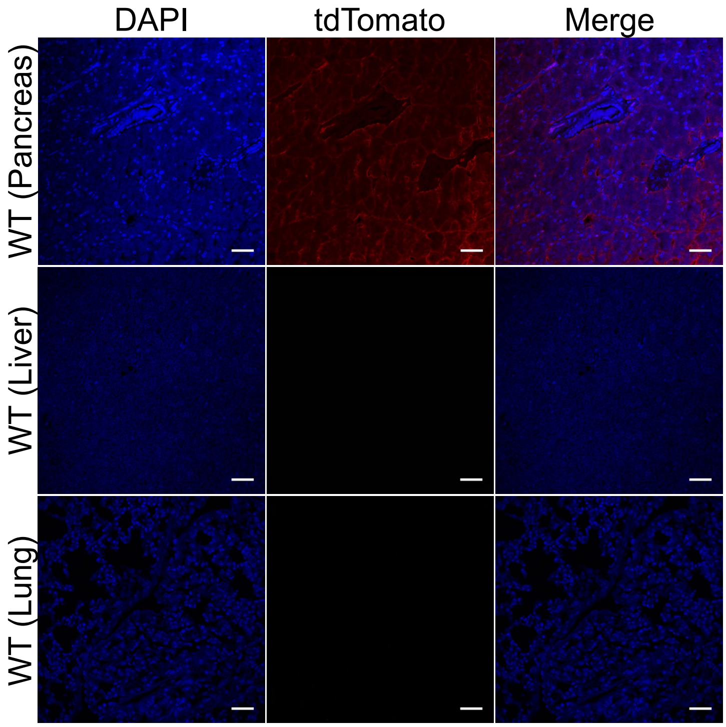
DAPI
tdTomato
Merge
WT (Pancreas)
WT (Liver)
WT (Lung)

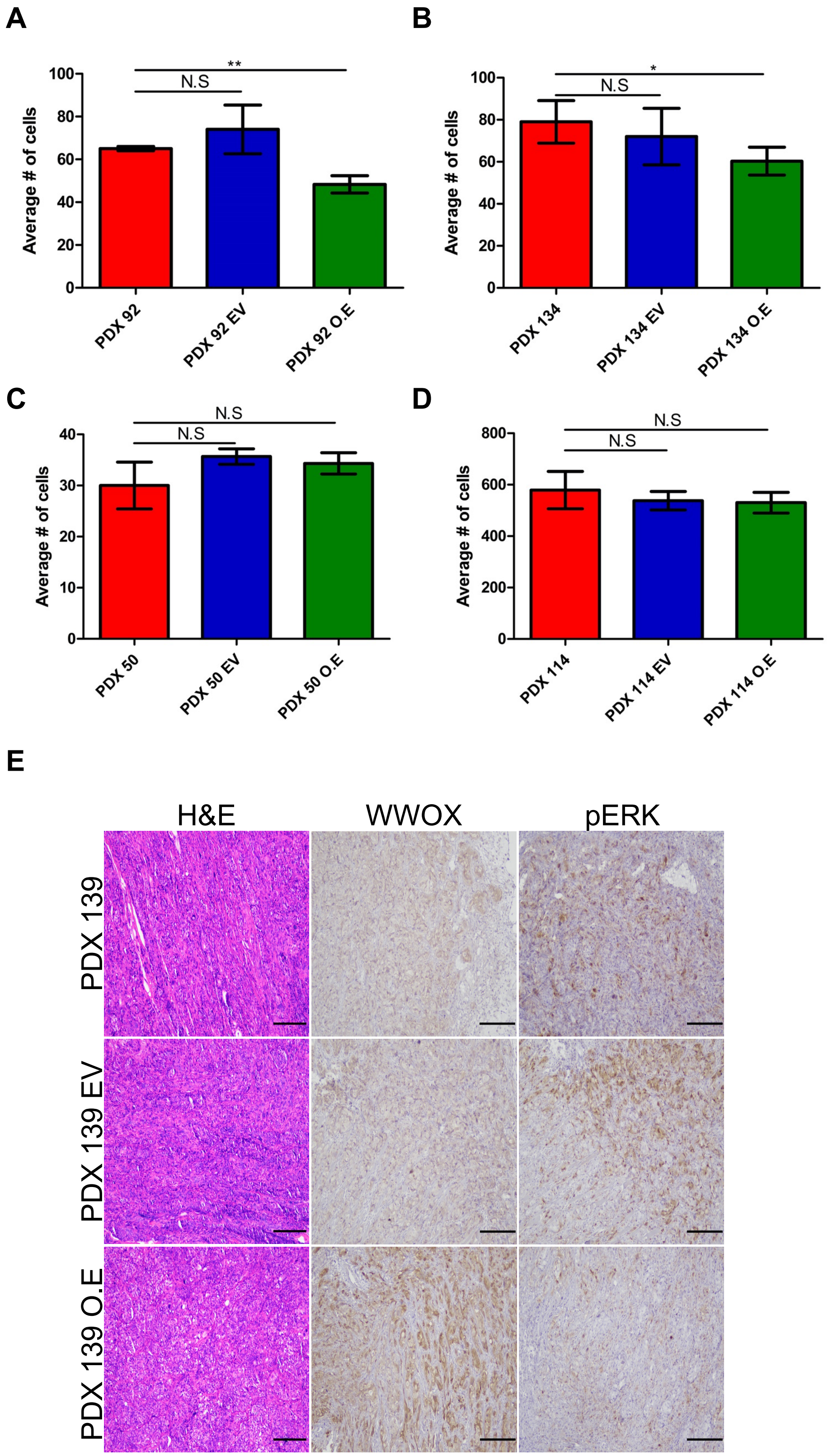

A
B
C
D
E
Average # of cells
N.S
**
*
PDX 92
PDX 92 EV
PDX 92 O.E
PDX 134
PDX 134 EV
PDX 134 O.E
PDX 50
PDX 50 EV
PDX 50 O.E
PDX 114
PDX 114 EV
PDX 114 O.E
H&E
WWOX
pERK
PDX 139
PDX 139 EV
PDX 139 O.E

### Supplementary Figure Legends

**Supplementary Figure 1. Validation and characterisation of KWC mouse model.** (A) Immunofluorescence stain of Tomato in tamoxifen induced, and non-induced pancreas. (B) Immunoblotting of WWOX in KWC and KC mice one-month post-tamoxifen injection. (C) Immunochemical stain of WWOX in WT, WC, KC, KWC (hetero), and KWC (homo) respectively two-months post-tamoxifen injection (magnification, ×20). (D) Co-immunofluorescence stain of tomato and amylase in WT mice (magnification, ×40). (E) H&E and pERK staining, respectively, in WC mice (magnification, ×10). (F) Representative amylase and Ck-19 immunofluorescence staining of KC, and KWC mice (magnification, ×10). (G) Quantification of amylase intensity in KC and KWC mice. (H) Quantification of Ck-19 intensity in KC and KWC mice. (I) RNAseq analysis of *Runx1, Onecut2,* and *Foxq1* genes in WT, KC, and KWC mice at two-months post-tamoxifen injection. *$P$<0.05, **$P$<0. 001, ***$P$<0. 0001. Two-tail unpaired t-test. Quantification graphs are presenting the means ± SD.

**Supplementary Figure 2. Gradual loss of WWOX expression upon lesions formation.** (A) Representative WWOX immunostaining at normal and lesion sites of KC mice eight-months post-tamoxifen injection (magnification, ×40). (B) Quantification of WWOX average intensity at normal sites, ADM sites, early and advanced PanINs. NS>0.05, **$P$<0. 001, ***$P$<0. 0001. Two-tail unpaired t-test. Quantification graphs are presenting the means ± SD.

**Supplementary Figure 3.** RT-PCR of *Ck-19*, *Fibronectin*, *Slug*, *Runx2* and *Angptl4* respectively of WT, KC, and KWC mice one-month post-tamoxifen injection.

**Supplementary Figure 4. WWOX deletion induces ERK MAPK signaling through TGFβ and BMP2 signaling pathway.** (A) Immunoblotting of WWOX, pERK and ERK of WT (n=2), KC (n=3), KWC *Wwoxf/+* (n=3), and KWC *Wwoxf/f* (n=2) mice at one-month post-tamoxifen injection. Anti-WWOX blot was run separately on a different gel, but same samples. (B) RNAseq analysis of *Angptl4* of WT, KC, and KWC mice two-months post-tamoxifen injection. (C) RNAseq analysis of *Bmp2* of WT, KC, and KWC mice two-months post-tamoxifen injection. (D) Representative IHC staining of pERK of KC, and KWC mice one-month post-tamoxifen injection (magnification, ×40). (E) Quantification of pERK positive cells in KC (n=4), and KWC (n=9) mice. NS>0.05, **P*<0.05, ***P*<0. 001. Two-tail unpaired t-test. Quantification graphs are presenting the means ± SD.

**Supplementary Figure 5. Validation of metastasis origin.** (A) 3 out of the 16 KWC tumor-bearing mice formed metastasis. (B) Representative WWOX, Pancytokeratin, pERK, pSTAT3, and amylase immunochemical staining respectively in normal lungs of KWC mice (magnification, ×10). (C) Representative immunofluorescence staining of tdTomato of WT pancreas, liver, and lung respectively (magnification, ×40).

**Supplementary Figure 6. WWOX restoration suppress growth.** (A) Quantification of colony formation of PDX 92, EV, and WWOX O.E, respectively. (B) Quantification of colony formation of PDX 134, EV, and WWOX O.E, respectively. (C) Quantification of colony formation of PDX 50, EV, and WWOX O.E, respectively. (D) Quantification of colony formation of PDX 114, EV, and WWOX O.E, respectively. (E) Representative H&E, WWOX, and pERK stain respectively of PDX 139, PDX 139 EV, and PDX 139 WWOX OE (magnification ×10). NS>0.05, **P*<0.05, ***P*<0. 001. Two-tail unpaired t-test. Quantification graphs are presenting the means ± SD.
